# Characterizing the allele- and haplotype-specific copy number landscape of cancer genomes at single-cell resolution with CHISEL

**DOI:** 10.1101/837195

**Authors:** Simone Zaccaria, Benjamin J. Raphael

## Abstract

Single-cell barcoding technologies have recently been used to perform whole-genome sequencing of thousands of individual cells in parallel. These technologies provide the opportunity to characterize genomic heterogeneity at single-cell resolution, but their extremely low sequencing coverage (<0.05X per cell) has thus far restricted their use to identification of the *total copy number* of large multi-megabase segments in individual cells. However, total copy numbers do not distinguish between the two homologous chromosomes in humans, and thus provide a limited view of tumor heterogeneity and evolution missing important events such as copy-neutral loss-of-heterozygosity (LOH). We introduce CHISEL, the first method to infer allele- and haplotype-specific copy numbers in single cells and subpopulations of cells by aggregating sparse signal across thousands of individual cells. We applied CHISEL to 10 single-cell sequencing datasets from 2 breast cancer patients, each dataset containing ≈2000 cells. We identified extensive allele-specific copy-number aberrations (CNAs) in these samples including copy-neutral LOH, whole-genome duplications (WGDs), and mirrored-subclonal CNAs in subpopulations of cells. These allele-specific CNAs alter the copy number of genomic regions containing well-known breast cancer genes including *TP53, BRCA2*, and *PTEN* but are invisible to total copy number analysis. We utilized CHISEL’s allele- and haplotype-specific copy numbers to derive a more refined reconstruction of tumor evolution: timing allele-specific CNAs before and after WGDs, identifying low-frequency subclones distinguished by unique CNAs, and uncovering evidence of convergent evolution. This reconstruction is supported by orthogonal analysis of somatic single-nucleotide variants (SNVs) obtained by pooling barcoded reads across clones defined by CHISEL.

## 1 Introduction

Single-cell DNA sequencing is a promising technology to quantify tumor heterogeneity and evolution with unprecedented resolution, enabling the identification of rare subpopulations of cells with distinct mutations and the inference of the evolutionary dynamics of cancer^1–4^. Recently, single-cell barcoding technologies, including Chromium Single Cell CNV Solution from 10X Genomics^5,6^ and direct library preparation^7,8^, have been used to perform low-coverage whole-genome sequencing of thousands of individual cells in parallel, overcoming the limited number of cells and the amplification/coverage biases of previous techniques^4^. Due to technical and financial limitations, these technologies have extremely low sequencing coverage (<0.05X per cell) which has thus far limited their application to the detection of large (≈3-5Mb) copy-number aberrations (CNAs) in individual cells. CNAs alter the number of copies of genomic regions, are frequent somatic mutations that drive cancer development^9–12^, play a crucial role in cancer treatment and prognosis^13,14^, and provide important markers for reconstruction of cancer evolution^15–19^.

Since the human genome is diploid, each CNA affects one *allele* of a genomic region located on either of the two homologous chromosomes (maternal and paternal), called *haplotypes*. Many methods have been developed to identify *allele-specific copy numbers*, which indicate the number of copies of each homolog, from bulk tumor sequencing data^20–32^. Moreover, multiple cancer studies have demonstrated the importance of deriving allele-specifc copy numbers^10,33–35^. For example, *copy-neutral loss of heterozygosity* (LOH) – where one allele is lost and the other duplicated so the total copy number remains 2 – is common in many cancers^33,36–41^. Allele-specific copy numbers have also been shown to be essential for accurate inference of *whole-genome duplications (WGDs)*^24,42^ and timing WGDs relative to other CNAs^10,24,43^.

Despite the demonstrated importance of allele-specific copy numbers, previous single-cell sequencing studies have assumed that low-coverage data is too shallow to obtain allele-specific information from single cells^7,8,44–46^. Existing methods for identifying CNAs from single-cell sequencing data^6–8,45–50^ are limited to the inference of *total copy number*, which indicates only the sum of copy numbers at each locus, by analyzing differences between the observed and expected number of sequencing reads aligned to a locus, or the *read-depth ratio (RDR)*. The signal to detect allele-specific copy numbers is the *B-allele frequency* (BAF), or relative proportions of reads from the two alleles of a genomic region; however, standard methods to calculate the BAF from individual germline heterozygous SNPs do not work with extremely low coverage sequencing data.

We introduce <u>C</u>opy-number <u>H</u>aplotype <u>I</u>nference in <u>S</u>ingle-cells using <u>E</u>volutionary <u>L</u>inks (CHISEL), the first method that infers allele-specific and haplotype-specific copy numbers in single cells from low-coverage DNA sequencing data. CHISEL amplifies the extremely weak signal in individual SNPs into a sufficiently strong signal to compute BAFs in genomic regions of modest size (≈5Mb) by combining reference-based phasing methods with a novel algorithm to phase short haplotype blocks across cells. CHISEL further phases allele-specific copy numbers across cells using an evolutionary model to derive *haplotype-specific copy numbers* that indicate the number of copies of the alleles located on the same haplotype in individual cells. CHISEL includes several other innovative features including global clustering of RDRs and BAFs along the *whole* genome and across *all* cells, and is the first method that includes BAF in the challenging inference of the genome ploidy of individual cells.

We applied CHISEL to analyze 10 single-cell datasets from 2 breast cancer patients, each dataset containing ≈2000 cells. CHISEL identified extensive allele-specific CNAs in these samples, including copy-neutral LOH, WGDs, and *mirrored-subclonal CNAs*. These latter events are haplotype-specific CNAs that alter the number of copies of the two distinct alleles of the same genomic region in different cells^51^. We used the haplotype-specific copy numbers derived by CHISEL to reconstruct a more refined and accurate view of tumor heterogeneity than previous total copy number analysis. We identified events that alter the copy number of well-known breast cancer genes and characterize key mechanisms in tumor progression, including potential precursors of WGDs and evidence of convergent evolution. Finally, we identified somatic single-nucleotide variants (SNVs) in subpopulations of cells with distinct haplotype-specific copy-number profiles by leveraging the pseudo-bulk nature of barcoded single-cell sequencing data. These SNVs provide orthogonal evidence for the phylogeny inferred by CHISEL compared to the phylogeny inferred by total copy numbers. The observed frequencies of SNVs as well as the distribution of clones across sections of the same tumor provides further support for the CHISEL phylogeny. CHISEL provides a powerful tool to realize the potential of single-cell whole genome sequencing for studies of tumor heterogeneity and evolution.

## 2 Results

### 2.1 CHISEL Algorithm

We developed the CHISEL algorithm to identify allele- and haplotype-specific copy-number aberrations (CNAs) from low-coverage single-cell DNA sequencing data (Fig. 1). CHISEL leverages information across hundreds-thousands of individual cells to overcome the low sequencing coverage (<0.05X) per cell. As in DNA sequencing of bulk samples^20–32^, CHISEL uses two quantities derived from aligned reads to estimate the number of copies, *ĉ_t_* and *č_t_*, of the two alleles of each genomic region *t*. The first quantity, the *read-depth ratio (RDR) x_t_*, is directly proportional to the total copy number *c_t_* = *ĉ_t_* + *č_t_*. The second quantity, the *B-allele frequency (BAF) y_t_*, measures the imbalance between the number of copies of the two alleles and corresponds to either 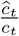 or 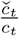. The key steps of CHISEL are: (1) computation of RDR and BAF in low-coverage DNA sequencing data from individual cells; (2) global clustering of RDRs and BAFs into a small number of copy-number states jointly across the *entire* genome of *all* cells; (3) inference of the pair {*ĉ_t_, č_t_*} of allele-specific copy numbers accounting for varying genome ploidy; (4) inference of haplotype-specific copy numbers (*a_t_, b_t_*) by phasing allele-specific copy numbers {ĉ_t_, č_t_}to their corresponding haplotypes across all cells; (5) inference of tumor clones by clustering of haplotype-specific copy numbers. We briefly describe these steps below and provide additional details in Methods.

**Fig. 1:**
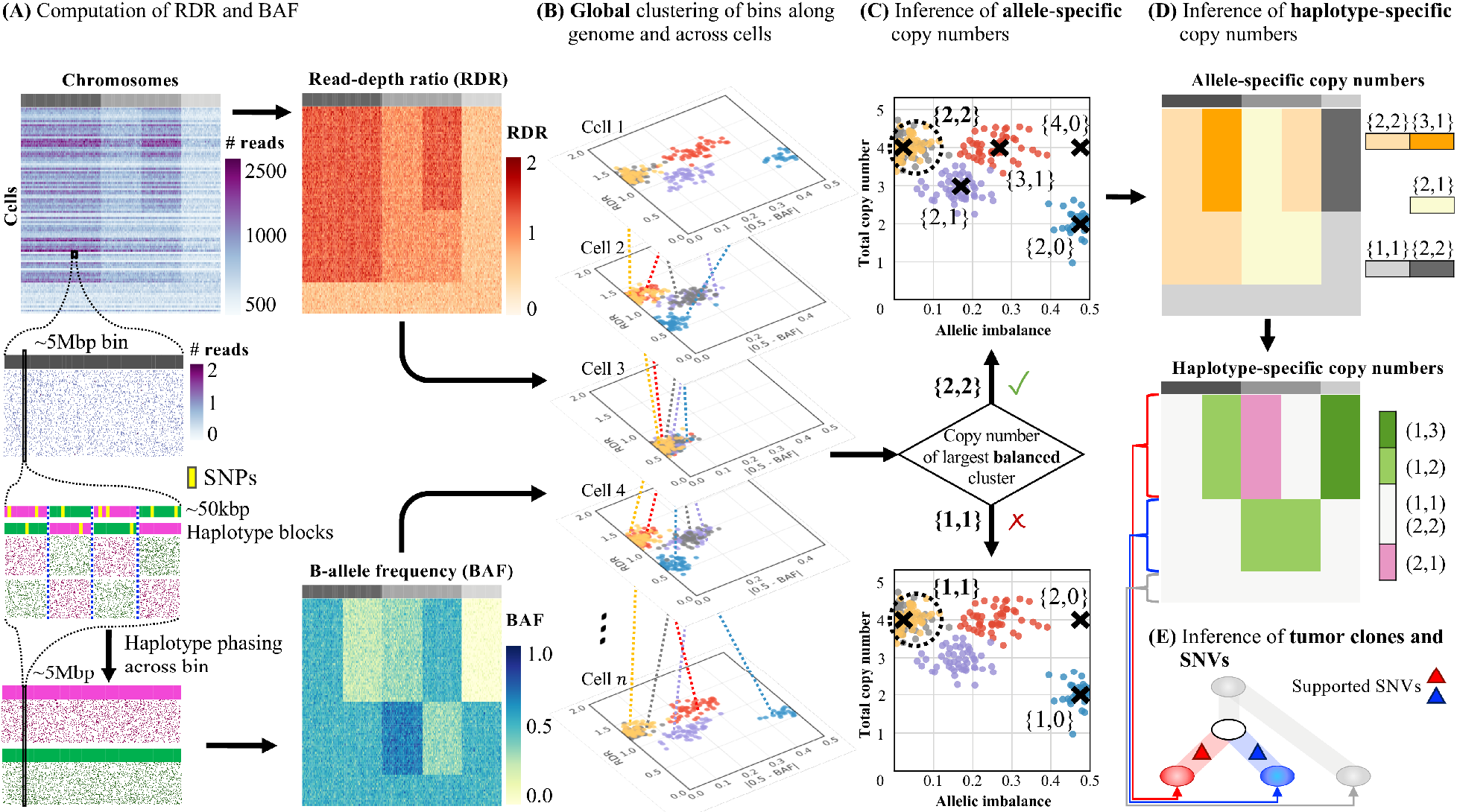
The CHISEL algorithm. **(A)** CHISEL computes RDRs and BAFs in low-coverage (<0.05X per cell) single-cell DNA sequencing data (top left). Read counts from 2 000 individual cells (rows) in 5Mb genomic bins (columns) across three chromosomes (grey rectangles in first row) are shown. For each bin in each cell, CHISEL computes the RDR (top) by normalizing the observed read counts. CHISEL computes the BAF in each bin and cell (bottom) by first performing referenced-based phasing of germline SNPs in ≈50kb haplotype blocks (magenta and green) and then phasing all these blocks jointly across all cells. **(B)** CHISEL clusters RDRs and BAFs *globally* along the genome and jointly across all cells resulting here in 5 clusters of genomic bins (red, blue, purple, yellow, and grey) with distinct copy-number states. **(C)** CHISEL infers a pair {ĉ, č}of allele-specific copy numbers for each cluster by determining whether the allele-specific copy numbers of the largest *balanced* (BAF ≈0.5) cluster are equal to {1, 1}(diploid), {2, 2}(tetraploid), or are higher ploidy. **(D)** CHISEL infers haplotype-specific copy numbers (a, b) by phasing the allele-specific copy numbers {ĉ, č}consistently across all cells. **(E)** CHISEL clusters tumor cells into clones according to their haplotype-specific copy numbers. Here, a diploid clone (light gray) and two tumor clones (red and blue) are obtained. A phylogenetic tree describes the evolution of these clones. Somatic single-nucleotide variants (SNVs) are derived from pseudo-bulk samples and placed on the branches of the tree.

In steps (1)-(3), CHISEL infers allele-specific copy numbers in individual cells. First, CHISEL divides the genome into bins of fixed size (here 5Mb). For each bin *t* in every cell, CHISEL computes the RDR *x_t_* using a standard normalization of the number of reads that aligned to *t*. Next, CHISEL computes the BAF *y_t_* by first using reference-based phasing algorithms^52^ to aggregate the individual SNPs in bin *t* into 50kb haplotype blocks and then phasing these blocks into the two alleles of *t* jointly across *all* cells (Fig. 1A). CHISEL is the first algorithm that calculates BAF in low-coverage single-cell DNA sequencing data. In Step (2), CHISEL globally clusters RDRs *x_t,i_* and BAFs *y_t,i_* into a small number of copy-number states across every bin *t* and cell *i* (Fig. 1B). This global clustering approach extends the one introduced in HATCHet^30^ for multi-sample bulk sequencing data. The global clustering leverages information along the entire genome and across all cells; in contrast current methods^6–8,45–50^ locally cluster RDRs of neighboring bins into segments and, with one recent exception^50^, analyze each cell independently. Finally in step (3), CHISEL infers the pair {*ĉ_t_, č_t_*} of allele-specific copy numbers for each bin t by identifying the largest balanced (BAF ≈ 0.5) cluster and using a probabilistic model-selection criterion to select the copy numbers (Fig. 1C). In contrast, existing approaches rely on the inference of the genome ploidy (or equivalent factors) from only RDRs^6–8,45–50^ and apply restrictive assumptions to select among many equally-plausible solutions without utilizing BAFs; this may result in selecting copy numbers that contradict the underlying allelic balance/imbalance (Fig. S17)

In steps (4) and (5), CHISEL infers haplotype-specific copy numbers (*a_t_, b_t_*) for each bin t in individual cells and clusters cells into clones according to these copy numbers. In contrast to the *unordered* pair {ĉ_t_, *c_t_*} of allelespecific copy numbers, the *ordered* pair (*a_t_, b_t_*) of haplotype-specific copy numbers indicates the number of copies of the alleles on each of the two homologous chromosomes, or *haplotypes*, 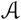 and 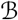. One cannot directly determine haplotype-specific copy numbers (*a_t_, b_t_*) of an individual cell from allele-specific copy numbers {*ĉ_t_, č_t_*} as we do not know the *phase* of each copy number, i.e. whether *a_t_* = *ĉ_t_, b_t_* = *č_t_* or *a_t_* = *c_t_, b_t_* = *ĉ_t_*. The key insight in deriving haplotype-specific copy numbers is to leverage the shared evolutionary history of the cells in a tumor and thus *phase* allele-specific copy numbers *jointly* across cells (Fig. 1D). CHISEL infers the phasing that minimizes the number of CNAs required to explain the haplotype-specific copy numbers using a model of interval events for CNA evolution^17–19^. This approach generalizes and extends the method introduced by Jamal-Hanjani et al.^51^ to infer haplotype-specific copy numbers in a restricted setting using multiple bulk-tumor samples. Finally, in step (5) CHISEL clusters cells into clones according to the inferred haplotype-specific copy numbers (Fig. 1E).

### 2.2 Allele-specific copy-number aberrations

We applied CHISEL to 10X Genomics Chromium single-cell DNA sequencing data from 2 breast cancer patients: patient S0 with 5 publicly available sequenced tumor sections and patient S1 with 5 previously-unpublished datasets. Each of the 10 datasets comprises ≈2 000 cells that were sequenced with coverage ranging from 0.02X to 0.05X per cell. CHISEL identified extensive allele-specific CNAs that were previously uncharacterized by total copy number analysis. Across all the datasets, we found that allele-specific CNAs alter ≈25% of the genome on average in at least 100 cells (Fig. S1). In addition, the novel features of CHISEL further improved the inference of total copy numbers (Supplementary Note B.1).

In patient S0, CHISEL identified 5-6 clones in each section that together comprise 70-92% of cells (Fig.S2-S6). In patient S1, CHISEL identified 2-3 clones in each section that together comprise 81-93% of cells (Fig. S1). For example, in section E of patient S0, CHISEL assigned 1448/2075 of cells to 6 clones, including one diploid clone (labeled I) and 5 aneuploid clones (labeled `II` - `VI`) (Fig. 2A). Since a single diploid clone and one or more aneuploid clones were identified in each tumor section, we concluded that the diploid clone comprises mostly normal cells while the aneuploid clones comprise tumor cells, in concordance with previous analysis^5^. The remaining 7-30% of cells are unclassified and the proportions of such cells are consistent with previously reported causes^5,6^ including cells in S-phase of the cell cycle with actively replicating DNA (12-42%), cells with a low number of sequenced reads (≈8%), and doublets (>2%). Interestingly, we found direct evidence of doublets in a small number of cells of patient S1 (Fig. S7).

**Fig. 2:**
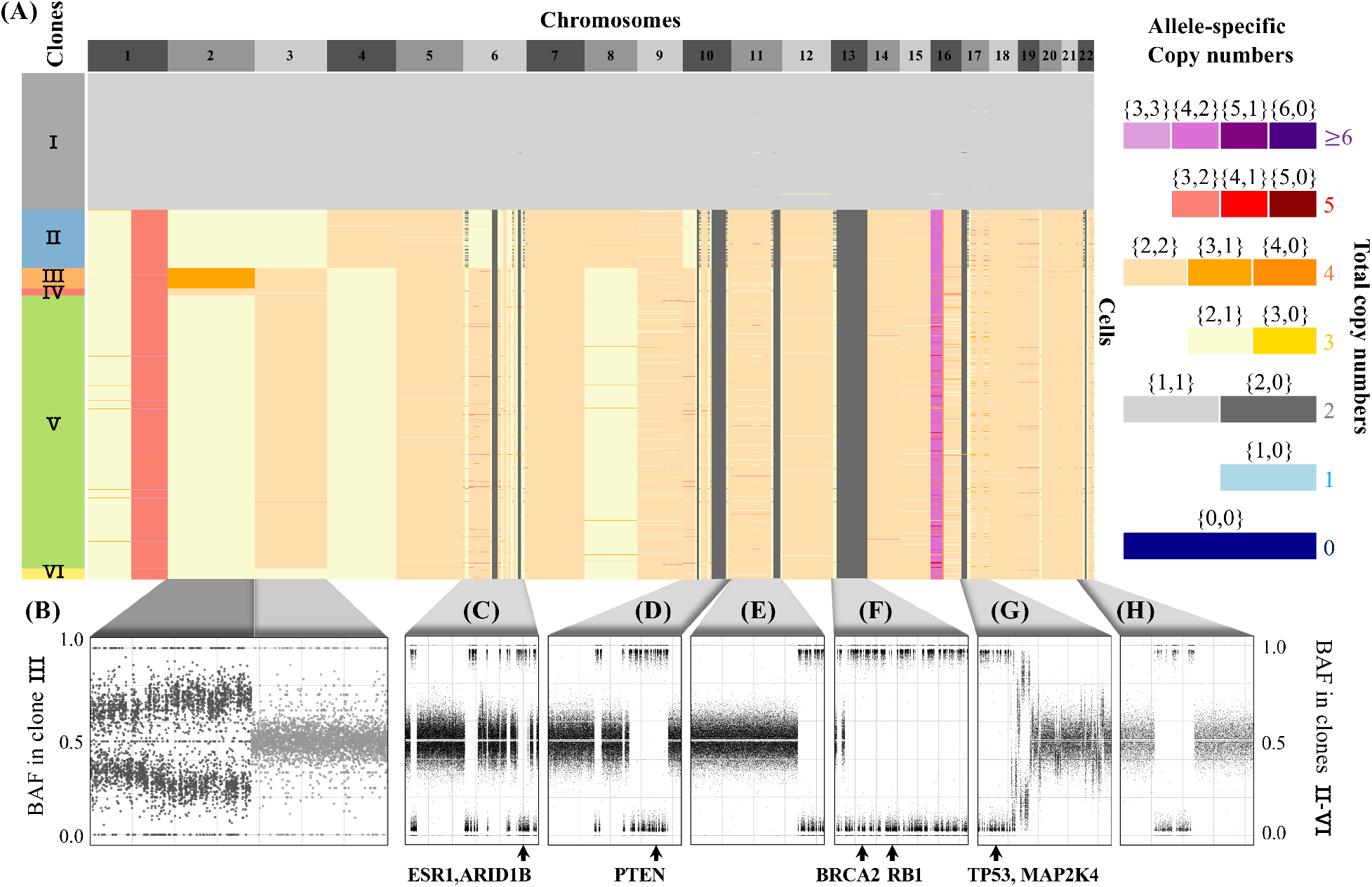
CHISEL reliably identifies allele-specific copy numbers. **(A)** Allele-specific copy numbers inferred by CHISEL across 1448 cells from section E of breast cancer patient S0. Clustering of cells according to allele-specific copy numbers reveals 6 clones (colored boxes on left). Clones `III` and `IV` are distinguished by an allele-specific CNA affecting the entire chromosome 2; cells in both these clones have 4 total copies of chromosome 2 but distinct allele-specific copy numbers equal to {3,1}and {2, 2}, respectively. **(B)** The BAF (computed in 50kb bins) of cells from clone `III` exhibits a clear shift away from BAF=0.5 in chromosome 2, supporting the unequal copy numbers of the two alleles on this chromosome. In contrast, chromosome 3 shows BAF ≈0.5, supporting equal copy numbers of the two alleles. **(C-H)** Copy-neutral LOHs are the most frequent allele-specific CNAs identified by CHISEL with examples shown on six chromosomes. All of these regions have a total copy number equal to 2, the same as the cells in the diploid clone I, but have allele-specific copy numbers equal to {2, 0}. Each of these copy-neutral LOHs is supported by BAFs approximately equal to 0 or 1 indicating the complete loss of one allele across all tumors cells. Known breast-cancer genes in each LOH region are indicated below each plot.

We found that allele-specific CNAs were important for resolving the clonal organization of the tumor. For example, in section E of patient S0, allele-specific copy numbers on chromosome 2 distinguish clones `III` and `IV` (Fig. 2A). Chromosome 2 has the same total copy number 4 in the cells of both clones `III` and `IV` but different allele-specific copy numbers of {3, 1}and {2, 2}, respectively. Tumor clones `III` and `IV` are thus indistinguishable using total copy numbers (Fig. S8). We found that BAFs support these allele-specific copy numbers with a significant shift above and below 0.5 observed along the entire chromosome 2 in the cells of clone `III` (Fig. 2B). The same allele-specific CNA is also observed in other sections from the same patient (Figs. S2-S5).

The most common type of allele-specific CNAs identified by CHISEL in both patients S0 and S1 are *copy-neutral LOHs* (Fig. S1), which have allele-specific copy numbers equal to {2, 0}. Copy-neutral LOHs are invisible to total copy number analysis methods as they are indistinguishable from normal diploid regions of the genome. We found copy-neutral LOHs in regions containing multiple genes implicated in breast cancer^53^ in patient S0 including *ESR1* and *ARID1B* on chromosome 6q, *PTEN* on chromosome 10q, *BRCA2* and *RB1* on chromosome 13, and *MAP2K4* on chromosome 17p (Fig. 2C-H). We also found copy-neutral LOHs in regions containing the genes *ESR1* and *ARID1B* on chromosome 6q in patient S1. Notably, CHISEL identified most of these copy-neutral LOHs to be clonal as they are present in nearly all tumor cells, suggesting an early acquisition of these mutations during the tumor evolution. These clonal copy-neutral LOHs are strongly supported by BAFs which clearly show the presence of a single allele in these regions across all tumor cells (Fig. 2C-H).

### 2.3 Allele and haplotype-specific mechanisms of tumor evolution

CHISEL derives *haplotype-specific copy numbers* in individual cells by examining changes in allele-specific copy numbers across cells. A particularly interesting application of haplotype-specific copy numbers is the identification of *mirrored-subclonal CNAs*: these are haplotype-specific CNAs occurring in different subpopulations of cells and affecting the two distinct alleles of the same genomic region. Such events were previously identified in the TRACERx multi-region sequencing of non-small-cell lung cancer patients and hypothesized to indicate parallel, or convergent, evolution^51^. We identified mirrored-subclonal CNAs on chromosomes 2 and 3 in a significant number of cells of patient S0 (Fig. 3A). Specifically, in section E we identified 168 cells with haplotype-specific copy numbers (1, 2) on chromosome 2 and 812 cells with haplotype-specific copy numbers (2, 1). To confirm the presence of this mirrored-subclonal CNA, we pooled sequencing reads from cells with the same haplotype-specific copy numbers and calculated the BAF in these two pseudo-bulk samples. The pooled BAFs across chromosome 2 show a clear switch in frequencies of the two haplotypes in the two subpopulations of cells (Fig. 3B). We observed the same mirrored-subclonal CNA in a large number of cells from the other sections of the same patient S0 (Fig. S2-S5). We note that multiple breast cancer tumor suppressor genes^53^ are present on these chromosomes, including *CASP8, MSH2*, and *DNMT3A* on chromosome 2, and *ATR, FOXP1*, and *SETD2* on chromosome 3. Therefore, mirrored-subclonal CNAs on these chromosomes suggest convergent evolution, as previously seen in non-small cell lung cancer^51^. We emphasize that mirrored-subclonal CNAs are not apparent to methods that calculate only total copy numbers or allele-specific copy numbers, and thus CHISEL’s ability to identify haplotype-specific copy numbers provides a refined view of the evolution of this tumor.

**Fig. 3:**
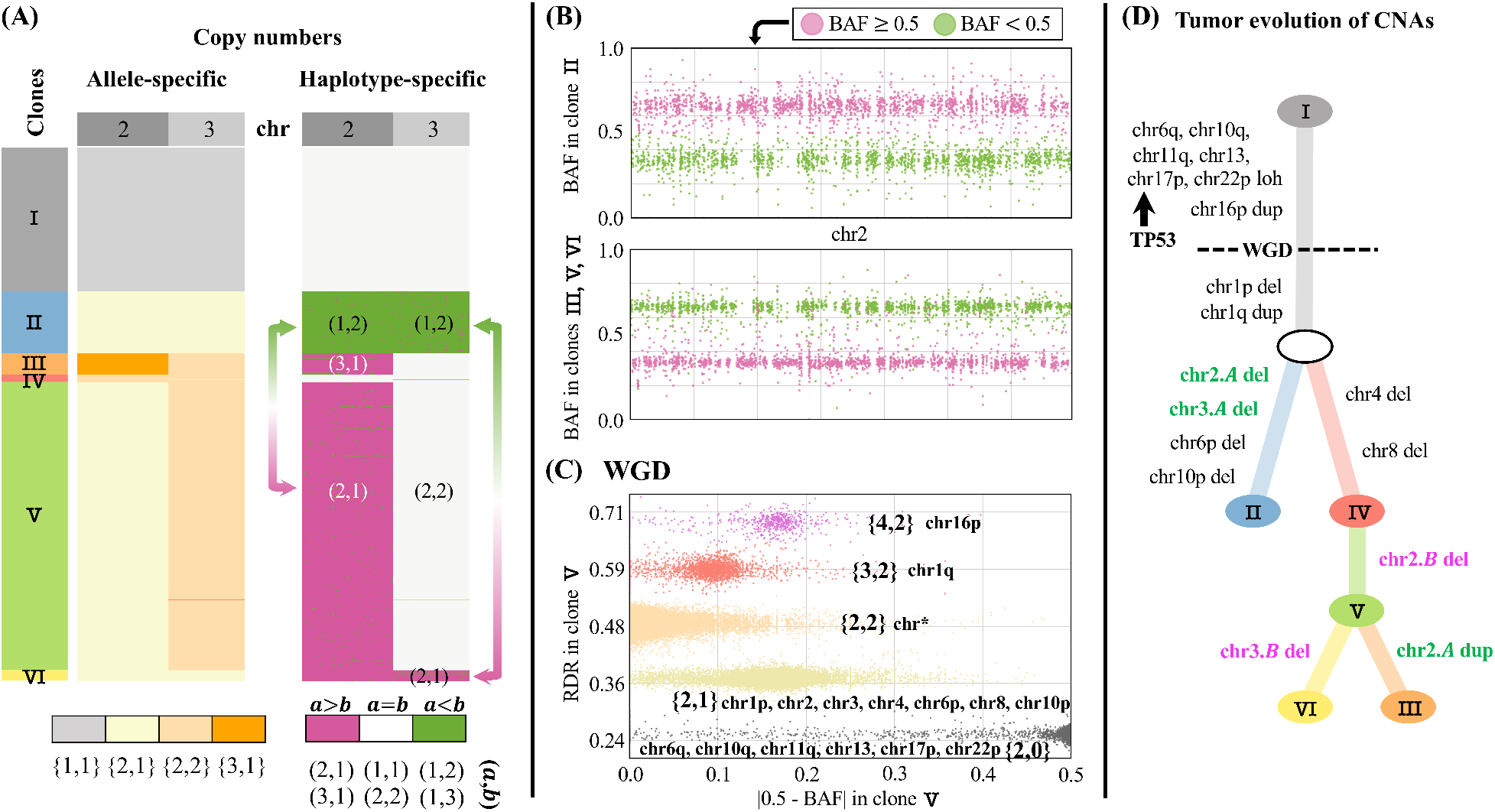
CHISEL reveals haplotype-specific CNAs and WGDs that shape tumor evolution. **(A)** CHISEL transforms allele-specific copy numbers (left) into haplotype-specific copy numbers (right) in 1448 cells from a breast tumor (section E of patient S0). Haplotype-specific copy numbers reveal *mirrored-subclonal CNAs* (arrows), or CNAs that alter the two distinct alleles of the same genomic region in different cells. Here, clone `II` has haplotype-specific copy numbers (1, 2) on chromosome 2, while clones `V` and `VI` have haplotype-specific copy numbers (2, 1). Similarly, clone `II` has haplotype-specific copy numbers (1, 2) on chromosome 3, while clone `VI` has (2, 1). **(B)** BAFs on chromosome 2 support mirrored subclonal CNAs, with a switch in the haplotype with larger BAF between clone `II` and clones `III`, `V`, and `VI`; each point in plot indicates BAF in a 50kb haplotype block. **(C)** RDRs, BAFs, and allele-specific copy numbers inferred by CHISEL along the entire genome and across all cells of clone `V` support the occurrence of a whole-genome duplication (WGD) as the two standard criteria for WGD are met: the the larger allele-specific copy number is greater than 2 in >50%of the genome and most of the genome (chr*) has allele-specific copy numbers {2, 2}. **(D)** A phylogenetic tree describes the CNA evolution of the 6 clones identified by CHISEL, with inferred haplotype-specific copy number events indicated on branches. LOH on multiple chromosomes and duplication of chromosome 16p precede the WGD, while deletion of chromosome 1p and duplication of chromosome 1q occur after WGD. Chromosome 17p contains the gene *TP53;* LOH at this locus supports published reports that *TP53* inactivation precedes WGD. Mirrored-subclonal CNAs on chromosomes 2 and 3 separate the 6 clones into two clear evolutionary branches, one containing the deletions of one haplotype of chromosomes 2 and 3 and the other containing the deletions of the other haplotype. These branches are further supported by the presence of specific subclonal CNAs, affecting chromosomes 4, 6p, 8, and 10p.

By integrating allele-specific copy numbers across cells, we inferred that a whole-genome duplication (WGD) was a clonal event that doubled the entire genome content of nearly all tumor cells in patient S0. We identified the occurrence of a WGD in every tumor clone of section E using criteria from published studies^24,42^ which demonstrated that allele-specific copy numbers are necessary to accurately infer the occurrence of WGDs. For example, the signal of WGD is clearly shown by pooling reads from all tumor cells of the largest tumor clone V of section E into a pseudobulk sample, and observing two imbalanced allele-specific deletion states, {2, 1}and {2, 0}(Fig. 3C). We similarly inferred the presence of a WGD in nearly all tumor clones from the other sections of patient S0 (Fig.S2-S5). In contrast, in patient S1 the inferred allele-specific copy numbers do not support the occurrence of a WGD and suggest that this tumor is mostly diploid (Fig. S1).

Using the haplotype-specific copy numbers, mirrored-subclonal CNAs, and WGDs identified by CHISEL, we constructed a phylogenetic tree that describes the tumor evolution of all 1448 cells in section E of patients S0 (Fig. 3D). The WGD and nine clonal CNAs that are present in all tumor cells are placed on the trunk of the phylogeny. The haplotype-specific copy numbers inferred by CHISEL enable the inference of the temporal order of some of these events^10,24^: the six clonal copy-neutral LOHs as well as the duplication of chromosome 16p are more likely to have occurred before WGD as the allele-specific copy numbers are even integers, while the two CNAs of chromosome 1 occurred after WGD. Interestingly, this inferred temporal order implies that LOH of chromosome 17p, which contains the gene *TP53*, *precedes* WGD, an order consistent with previous reports of *TP53* inactivation occurring before WGDs^42^. The mirrored-subclonal CNAs affecting chromosomes 2 and 3 separate the tumor clones into two clearly distinct branches: one including 168 cells from clone `II` and the other including 890 cells from the other clones. The two distinct branches are further subdivided by subclonal CNAs that are unique to each branch: CNAs of chromosomes 6p and 10p are unique to clone `II` while CNAs of chromosome 4 and chromosome 8 are unique to the other tumor clones.

### 2.4 Clonal evolution across multiple tumor regions and somatic single-nucleotide variants

We applied CHISEL to jointly analyze 10 202 cells from all 5 sequenced sections of breast cancer patient S0 (Fig. 4A). Using the inferred allele- and haplotype-specific copy numbers (Fig. S9), we constructed a phylogenetic tree describing the evolution of 8 tumor clones (labeled `J-I - J-VII`) that include 4 085 cells across all 5 sections (Fig. 4B) and one normal diploid clone that includes 4 239 cells. This tree recapitulates the major features of the tree inferred from only the cells in section E (Fig. 3D), including: clonal CNAs and copy-neutral LOHs that occur before and after a WGD, deletions on chromosomes 4, 6p, 8, and 10p that define the initial split into a *left* branch containing clones `J-I` and `J-II`, and a *right* branch containing clones `J-III - J-VIII`, and mirrored-subclonal CNAs on chromosomes 2 and 3 that further subdivide the subclones in these two branches. The larger number of cells in the integration of data from all 5 tumor sections yields a more refined tree with additional clones that contain small numbers of cells and are defined by haplotype-specific CNAs; e.g. clones `J-I, J-IV`, and `J-VIII` in Fig. 4B.

**Fig. 4:**
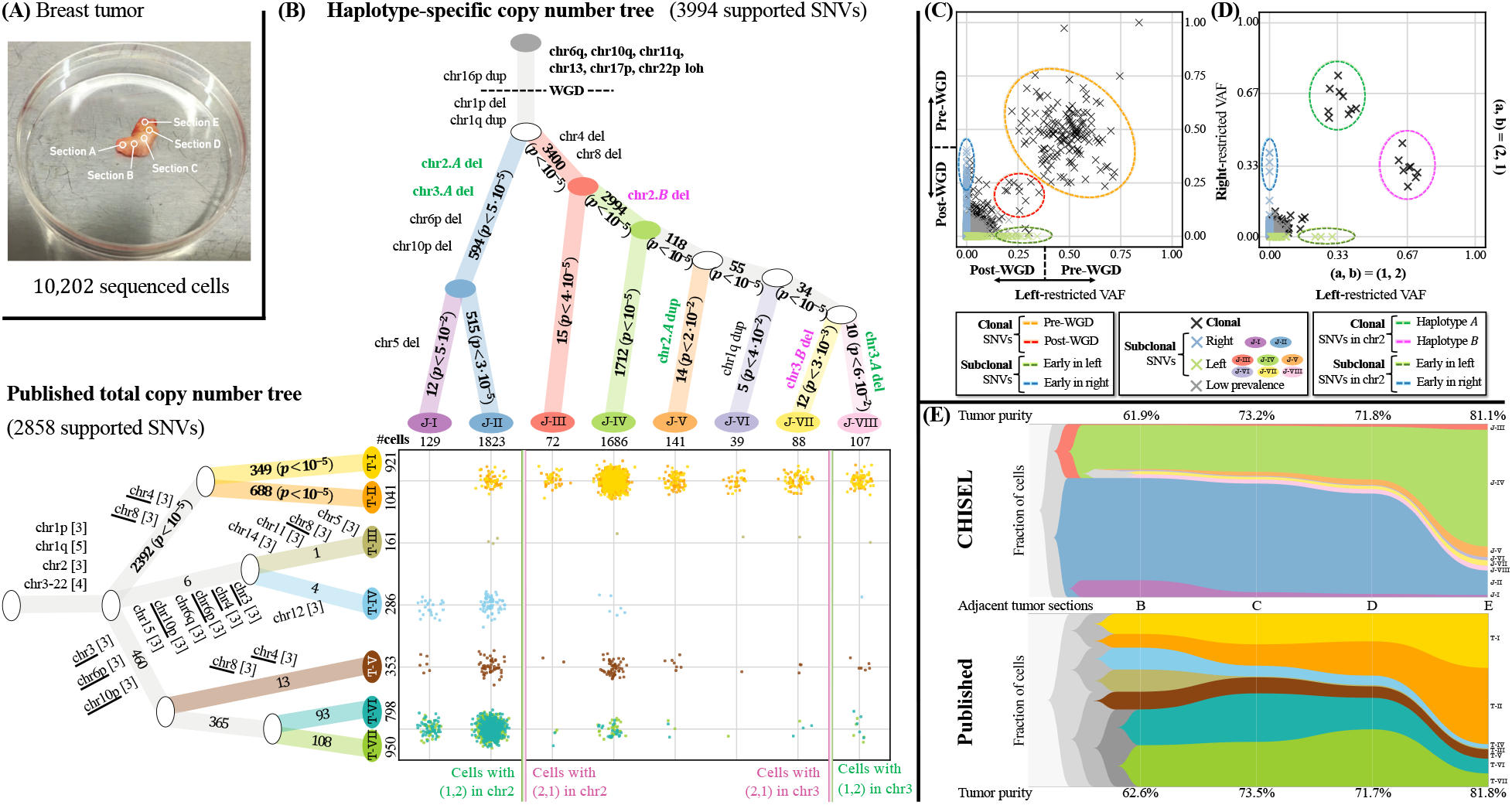
Reconstruction of tumor heterogeneity and evolution across multiple tumor sections. (**A**) DNA from 10 202 cells in five adjacent sections of a breast tumor patient S0 was sequenced using the 10X CNV Solution. (**B**) Phylogenetic trees describing tumor clones and their CNA evolution as inferred by CHISEL jointly across all cells (top) and as reported in previous analysis using total copy numbers (left). Each branch of CHISEL tree is labeled by haplotype-specific copy-number events, and each branch of the total copy number tree is labeled by total copy number changes (in brackets with repeated events underlined). Each non-truncal branch in the trees is also labelled by the number of supported SNVs; *p*-values comparing to the number of such SNVs expected by chance are indicated, with significant values (*p* < 10^−1^) in boldface. Overall, 3994 SNVs support the haplotype-specific tree inferred by CHISEL, but only 2858 SNVs support the published tree inferred using total copy numbers. (Middle) Comparison of the cellular composition of clones identified in the two analysis, where each point corresponds to a cell colored by its clone assignment. For simplicity, clones `T-I` and `T-II` as well as `T-VI` and `T-VII` are merged as they are distinguished by only few isolated and small (<5Mb) CNAs. Vertical lines separate cells with different haplotype-specific copy numbers in chromosomes 2 and 3 in the CHISEL tree. (**C**) Left-restricted VAF (resp. right-restricted VAF) is computed as the variant-allele frequency (VAF) of SNVs using sequencing reads from cells in either the left branch (clones `J-1` and `J-II`) (resp. right branch with clones `J-III - J-VIII`) of CHISEL tree. Excluding low-prevalence SNVs that are potential false positives, the CHISEL tree partitions SNVs according to their clonal status (clonal or subclonal in each branch) and the restricted VAFs support this classification. Clonal SNVs have restricted VAFs consistent with their occurrence before (≈0.5) and after (⩽0.25) WGD and all subclonal SNVs have restricted VAF consistent with their occurrence after WGD (≤0.25). Subclonal SNVs with the highest restricted VAFs (some ≈0.33 due to CNAs) are inferred to occur early in the evolution of these clones. (**D**) Restricted VAFs of all clonal SNVs on chromosome 2 occurring before WGD are consistent with the different haplotype-specific copy numbers (1, 2) and (2, 1) in the left and right branches: SNVs on the two distinct haplotypes have restricted VAFs equal to 0.33 and 0.67 in the two branches. Early SNVs in each branch have restricted VAFs ≈0.33, which is consistent with occurrence after WGD. (**E**) The proportions of tumor clones and the corresponding tumor purity identified by CHISEL and total copy-number analysis across the the four adjacent tumor sections with highest tumor purity.

We compared the CHISEL tree with a previously described tree^5^ derived from the total copy numbers (Fig. S10) inferred by Cell Ranger DNA^48^ (reported to be consistent^6^ with copy numbers obtained with Ginkgo^45^) and containing 7 tumor clones (labeled `T-I` - `T-VII`). We found that there is good agreement on the initial branch of both trees (Fig. 4B): deletions of chromosomes 4 and 8 occur on the branch containing clones `J-III - J-VIII` in the joint tree and clones `T-I` and `T-II` in the total copy number tree, while deletions of chromosomes 3, 6p, and 10p occur on the the branch containing the remaining clones in both trees. However, CHISEL further subdivided cells into novel clones/subpopulations that are characterized primarily by allele- and haplotype-specific CNAs. Of particular note are the mirrored-subclonal CNAs identified by CHISEL on chromosomes 2 and 3 that distribute cells from clones `T-I` and `T-II` in the tree derived from total copy numbers into clones `J-III` - `J-VIII` in the tree derived from CHISEL. These mirrored-subclonal CNAs are invisible to the total copy number analysis and consequently the tree constructed from total copy numbers includes cells with different haplotype-specific copy numbers in the same clone (e.g. `T-V`) and infers multiple independent occurrences of the same copy-number events on different branches of the tree (e.g. chromosomes 4, 6p, 8, and 10p) (Fig. 4B).

To further quantify the differences between the phylogenetic trees produced by CHISEL and total copy number analysis, we examined somatic single-nucleotide variants (SNVs). Since SNVs were not used in tree construction, they provide an orthogonal signal for subdividing cells into subpopulations. Because of extremely low sequencing coverage, identification of SNVs in individual cells is impossible. Thus, we pooled sequencing reads from all cells into a pseudo-bulk sample, and identified ≈49k SNVs using standard methods developed for bulk-tumor sequencing data^54^. We assigned each SNV to those cells with a variant read and found that 10 551 of the SNVs are present only in the tumor clones `J-1 - J-VIII`. This number of SNVs is close to the average of ≈7000 (range 500–93 000) somatic SNVs reported in whole-genome sequencing studies of 560 breast tumors^53^. Next, for each non-truncal branch in the phylogenetic trees, we computed the number of SNVs that are uniquely assigned to cells in the subtree defined by that branch. We found that ≈40%more SNVs (3994 vs. 2858) are consistent with the tree inferred by CHISEL compared to tree inferred by total copy numbers. Moreover, we found that all 14 branches in the CHISEL tree are supported by more SNVs than expected by chance (*p* < 10^−1^), while only 3/11 branches in the total copy number tree have significant support (Fig. 4B). Additionally, we found that while clones `T-I` and `T-II` in the total copy number tree are supported by a significant number of SNVs – even though they are not distinguished by any large CNAs (Fig. S10C) - these SNVs are the same as those that support the smaller subclones (`J-III - J-VII`) identified by CHISEL. In summary, we found that SNVs support nearly all tumor clones inferred by CHISEL in both patients S0 andS1 (Fig. S11 and Fig. S12).

Next, we examined the relationship between the variant-allele frequency (VAF) of each SNV – the proportion of reads covering an SNV locus that contain the variant allele – and the clonal status of the SNV induced by the CHISEL tree. We classified the SNVs according to the partition of cells defined by the initial two branches of the CHISEL tree and, after excluding likely false positive SNVs with low clone prevalence, we obtained: 594 SNVs unique to cells in the left branch (clones `J-I` and `J-II`), 1632 SNVs unique to the right branch (clones `J-III - J-VIII`), and 2 798 clonal SNVs in both branches. Since each read has unique cell barcode we computed the *left-restricted VAF (resp*. right-restricted VAF) of each SNV using only the sequencing reads from the subpopulation of cells in the left (resp. right) branch of the CHISEL tree. We found that restricted VAFs of SNVs were consistent with the placement of the SNVs on the CHISEL tree (Fig. 4C): clonal SNVs have restricted VAFs consistent with their occurrence before (≈0.5) or after (≈0.25) WGD (assuming no other CNAs at the locus) in *both* branches, while subclonal SNVs have lower restricted VAFs (⩽0.25). In addition, we found that the restricted VAFs of SNVs on chromosome 2 are consistent with the corresponding mirrored-subclonal CNA (Fig. 4D): clonal SNVs that occurred before WGD have restricted VAF equal to either ≈0.33or ≈0.67 when they are located on the deleted allele or the other allele, respectively. SNVs with both these values of restricted VAF in the two distinct branches clearly support the deletion of two different haplotypes. We observed similar consistency between the standard VAF computed across all cells and the placement of the SNV on the CHISEL tree (Fig. S13).

We observed an interesting discordance between the number of cells in the left (clones `J-I` and `J-II`) and right (clones `J-III - J-VIII`) branches and the number of SNVs assigned to these branches. While the left and right branches have a very similar number of cells (1952 vs. 2133, respectively), the left branch has fewer SNVs (594 vs. 1632). This discordance may reflect different rates of growth and/or selection between the clones in these branches. Intriguingly, we found a subclonal CNA affecting the entire HLA gene complex in chromosome 6p that is unique to the left branch and could provide a mechanisms for evasion of immune response^55^.

Finally, we examined the variation in proportion of cells in each tumor clone across the different sections of patient S0, as these are adjacent sections of the same tumor. We found that the left and right branches in the CHISEL tree are consistent with the spatial distribution of the tumor as *all* the clones in the same branch consistently expand or contract across the adjacent tumor sections B-E (Fig. 4E): clones `J-I` and `J-II` from the left branch contract towards section E, while all the remaining clones from the right branch expand towards section E. In contrast, the clones inferred by total copy number analysis have more complicated dynamics across the tumor sections (Fig. 4E). While the merge of clones `T-VI` and `T-VII` contract towards section E and the merge of clones `T-I` and `T-II` expand towards section E, both the proportions of the subclones in these groups and the proportions of the remaining clones fluctuate independently across the sections. This discordance between the spatial and temporal evolution suggests that the clones inferred by the total copy number analysis are less plausible.

## 3 Discussion

New technologies to perform low-coverage whole-genome sequencing from thousands of individual cells provide data to study tumor heterogeneity and evolution at previously unprecedented resolution. However, methods to analyze this data have thus far been limited to the identification of total copy numbers of large genomic regions in individual cells. Here, we introduce CHISEL, the first algorithm to infer allele- and haplotype-specific copy numbers in single cells from low-coverage DNA sequencing data. CHISEL integrates the weak allelic signals across thousands of individual cells, leveraging a strength of single-cell sequencing technologies (many cells) to overcome a weakness (low genomic coverage per cell). CHISEL also includes other innovative features, such as global clustering of RDRs and BAFs, and a rigorous model selection procedure for inferring genome ploidy, that improves both the inference of allele-specific and total copy numbers.

We demonstrated the unique features of CHISEL on 10 datasets, each comprising ≈2 000 cells, from 2 breast cancer patients. CHISEL identified previously-uncharacterized CNAs and mutational events that shape the tumor heterogeneity and evolution, including extensive allele-specific CNAs – especially copy-neutral LOHs – that further distinguish novel clones and affect well-know breast cancer genes. In addition, the allele- and haplotype-specific copy numbers inferred by CHISEL reveal mirrored-subclonal CNAs and whole-genome duplications (WGDs) that characterize some of the key mechanisms of tumor evolution, including evidence of convergent evolution and potential precursors of WGDs. Many of these events are corroborated by somatic single-nucleotide variants (SNVs) and the spatial distribution of the inferred clones. Overall, CHISEL provides a more refined and accurate view of tumor evolution than the one obtained by previous total copy number analysis.

The single-cell view of allele- and haplotype-specific copy numbers provided by CHISEL offers the opportunity for deeper analysis of tumor evolution. CHISEL enables the identification of allele-specific CNAs, including copy-neutral LOHs and whole-genome duplications (WGDs), and haplotype-specific CNAs, including mirrored-subclonal CNAs, in individual cells without the limitations of bulk tumor-samples where inference of tumor ploidy and purity from admixed signals is extremely challenging^20–32^. In addition, previous analysis of haplotype-specific CNAs have been restricted to the special case where these CNAs are present in different samples from the same patient^51^. CHISEL may be used to analyze the frequency and function of mirrored-subclonal CNAs and other complex copy number events across different cancer types, especially for haplotype-specific CNAs which have received scant attention thus far in analysis of bulk-tumor samples.

While CHISEL enables the accurate inference of allele- and haplotype-specific copy numbers in individual cells, there are a number of areas for future improvements. First, the low coverage of single-cell DNA sequencing data limits the size of the CNAs that can be accurately inferred. One approach to improve the resolution is to iteratively run CHISEL on pseudo-bulk genomes obtained by merging multiple cells with the similar haplotype-specific copy number profiles, as suggested in recent studies^7,8^. Second, BAF estimation could be further improved by using larger haplotype blocks that one could infer from the results of CHISEL in regions of allelic imbalance or using the signal from sequencing reads that cover multiple SNPs. Third, a more refined model of CNA evolution might be integrated in the inference of haplotype-specific copy numbers, for example reconstructing the full copy-number tree from the model of interval events^19^ or integrating the additional signal from breakpoints^32^. Fourth, one could improve techniques for classifying cells with highly aberrant copy-number profiles, designing classifiers to distinguish actively replicating regions^6,8^ from cell doublets.

CHISEL’s results also provide a useful substrate for other analyses of tumor heterogeneity and evolution. In particular, further integration of CNAs and SNVs in single-cells would provide higher resolution reconstructions of tumor evolution. While our initial analysis of SNVs using restricted-VAFs showed good consistency between SNVs and the clones inferred from CNAs, a complete and accurate classification of all SNVs remains a challenging problem: distinguishing true SNVs from false positive is difficult for clones with few cells, and also variant read counts are expected to be low in regions of high copy number (e.g. from WGD). The SNV analysis could be further extended to derive the mutant copy number of individual SNVs, a task that is notoriously difficult in bulk tumor sequencing where discordance between variant allele frequencies (VAFs) and cancer cell fractions (CCFs) complicates tumor evolution studies^56–59^. In addition, integrated analysis of CNA and SNV evolution would help resolve questions about the relative rates of these mutation classes during tumor evolution, including “punctuated evolution” of CNAs in certain tumors^60^. Our analysis of tumor patient S0 showed an intriguing discordance between number of SNVs unique to a copy number clone and the prevalance of the clone in the tumor cell population, and further studies of such phenomenon in additional tumors would be informative. Finally, allele-specific copy numbers provide useful signal for single-cell studies of allele-specific gene expression by combining single-cell DNA sequencing with single-cell RNA sequencing^44,61^.

## 4 Method

### 4.1 CHISEL Algorithm

We introduce the <u>C</u>opy-number <u>H</u>aplotype <u>I</u>nference in <u>S</u>ingle-cells using <u>E</u>volutionary <u>L</u>inks (CHISEL) algorithm to derive allele- and haplotype-specific copy numbers from low-coverage DNA sequencing of *n* cells by integrating two signals, the read-depth ratio (RDR) and the B-allele frequency (BAF), *jointly* across the whole genome of *all* cells. We divide the reference genome into *m bins* and represent the genome of each cell by two integer vectors, **a** = (*a*_1_,…, *a_m_*) and **b** = (*b*_1_,…, *b_m_*). Each bin t has two alleles, and the entry *a_t_* indicates the number of copies of the allele that is located on haplotype 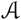 whereas the entry *b_t_* indicates the number of copies of the other allele located on haplotype 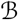. We call (**a**, **b**) the haplotype-specific copy numbers of the cell.

CHISEL addresses three major challenges in the derivation of the haplotype-specific copy numbers of each cell from low-coverage single-cell DNA sequencing data. The first is the calculation of the BAF as the extremely low-coverage data precludes the standard approach used in bulk sequencing that estimates the BAF from the proportion of alternate reads at individual germline SNPs^20–30^. The second is the inference of a pair {*ĉ_t_, č_t_*}of allele-specific copy numbers for each bin *t* in the presence of varying genome ploidy across cells resulting from CNAs and whole-genome duplications (WGDs). The third is the inference of an *ordered* pair (*a_t_, b_t_*) of haplotype-specific copy numbers from the *unordered* pair {*ĉ_t_, č_t_*}of allele-specific copy numbers, as we do not know *a prior* which allele is located on haplotype 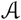 or 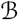.

CHISEL has five major steps (Fig. 1) which we detail in the subsections below.

#### 4.1.1 Computation of read-depth ratio and B-allele frequency

The first step of CHISEL is to compute the RDRs x_*i*_ = (*x*_1,*i*_,…, *x_m,i_*) and the BAFs *y_i_* = (*y*_1,*j*_,…, *y_m,i_*) for all m bins in every cell *i* (Fig. 1A). The RDR *x_t,i_* of bin *t* in cell *i* is directly proportional to the total number of reads that align to *t* and is used to estimate the total copy number *c_t,i_* = *č_t,i_* + *ĉ_t,i_*; i.e. *c_t,i_* ≈ *γ_i_x_t,i_* for some cell-specific scale factor *γ_i_*. CHISEL computes *x_t,i_* by appropriate normalization of the number of reads aligned to sufficiently large bins (5Mb in this work) – accounting for GC bias and other biases – similar to other approaches for CNA detection from bulk^20–32^ or single-cell^6–8,45–50^ sequencing (Supplementary Note C.1).

The BAF *y_t,i_* of bin *t* in cell *i* is the fraction of reads belonging to one of the two distinct alleles of *t* and provides an estimate of either 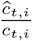 or 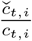. Previous methods for bulk sequencing data compute the BAF from either individual heterozygous germline SNPs^20–30^ or by aggregating SNPs in small haplotype blocks^31,32^. However, methods that compute BAF from individual SNPs are not useful for low-coverage DNA sequencing data because few, if any, reads will cover each SNP; e.g. in the 10X Genomics datasets that we analyzed (≈0.04X coverage) only ≈2% of SNPs were covered by at least one read in a single cell and only ≈0.08% were covered by more than 1 read. Methods that compute BAF from haplotype blocks are also not directly applicable since the inferred blocks remain too short, containing too few SNPs for accurate calculation of BAF in single cells. Current reference-based phasing methods have switch error rates^62^ of ≈1%meaning that haplotype blocks longer than a few hundred kilobases are likely to contain a phasing error. For example, the Battenberg algorithm^31^ for bulk DNA sequencing reported blocks up to ≈300kb, which is too small to contain enough SNP-covering reads for accurate calculation of BAF in individual cells.

CHISEL computes the BAF *y_t,i_* of bin t in cell i using a two-stage procedure. In the first stage, CHISEL uses the reference-based algorithm Eagle2^52^ to phase germline heterozygous SNPs in each bin *t* into *k* haplotype blocks; we use blocks of length 50kb in this work as phasing errors are unlikely at this scale^62^. Each haplotype block is composed of two sequences of nucleotides at consecutive SNPs, called the reference and alternate sequences, with each sequence located on a different allele of *t*. In the second stage, CHISEL computes the BAF *y_t,i_* by phasing the *k* blocks into the two alleles of bin *t* across *all* cells. Specifically, we phase the *k* blocks with respect to one of the two alleles of *t*, which we call the *minor allele* 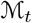(see below), and we define the *phase h_p_* of every block *p* such that *h_p_* = 1 indicates that the alternate sequence of p belongs to 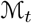, and *h_t_* = 0 otherwise. Given the phases *h*_1_,…, *h_k_* of the *k* blocks in bin *t*, CHISEL calculates the BAF *y_t,i_* from the total number *T_P,i_* of reads covering block *p* in cell *i* and the corresponding number *V_p,i_* of reads only covering the alternate sequence of p in cell i as follows

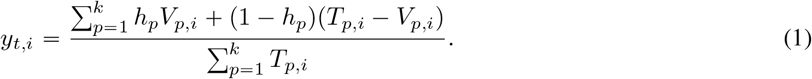

The BAF is typically calculated in bulk sequencing to estimate the proportion of an unknown allele of *t* and corresponds to either 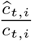 or 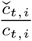. In contrast, CHISEL calculates *y_t,i_* to estimate the proportion of the *same* allele 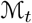 in every cell *i*. As the phases *h*_1_,…, *h_k_* are unknown, CHISEL seeks values of *h*_1_,…, *h_k_* such that *y_t,i_* in Eq. (1) accurately estimates 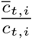 where 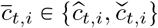 is the copy number of 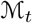 in every cell *i*.

CHISEL infers phases *h*_1_,…, *h_k_* for the blocks in bin *t* based on the estimated proportion *Y_t_* of the minor allele 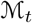 across all cells, where we assume without loss of generality that 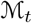 is the allele of *t* with the lower proportion so that 0 ⩽ *Y_t_* ⩽ 0.5. We designed an expectation-maximization (EM) algorithm^63^ to calculate the maximum likelihood estimate *Y_t_* given the observed values of read counts *T_P,i_* and *V_p,i_* in every block *p* across *every* cell *i*, where the phases *h*_1_,…, *h_k_* are unobserved, latent variables (Supplementary Note C.2). One issue that arises in this calculation is that the cases *Y_t_* ≉ 0.5 and *Y_t_* ≈ 0.5 need to be handled differently. When *Y_t_* ≉ 0.5 we compute the maximum-likelihood phases *h*_1_,…, *h_k_* (Theorem S1) using the idea that the lower (respectively higher) read counts belong to the same allele where there is allelic imbalance, as previously used in bulk sequencing^20,23,29^. We also showed empirically that this approach accurately identifies both the true phases and the BAF *y_t,i_*(Fig. S14). When *Y_t_* ≈ 0.5, the allelic origin of each block cannot be determined from read counts^29^. Thus, CHISEL selects the haplotype phase *h_p_* of every block p uniformly at random. We show that the corresponding *y_t,i_* is an unbiased estimator of 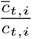 under the assumption that *Y_t_* ≈ 0.5 implies 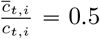 in every cell *i* (Theorem S2), which is reasonable as the only violations are rare. Further details are in Supplementary Note C.3.

#### 4.1.2 Global clustering of genomic bins into copy-number states

The second step of CHISEL is to cluster bins into a small number of copy-number states according to the RDR *x_t,i_* and BAF *y_t,i_* values for every bin *t* in cell *i*. Clustering helps to overcome measurement errors and variance in the computed RDRs and BAFs. The standard approach used in bulk^20–32^ and single-cell^6–8,45–50^ copy number analysis is to segment bins *locally* along the genome grouping neighboring bins with similar values of RDR and/or BAF into segments that are assigned the same copy-number state. This local segmentation leverages the observation that a CNA alters the copy numbers of multiple adjacent bins. Existing methods for single-cell copy numbers^6–8,45–50^ perform this segmentation on RDR values only as these methods do not calculate BAF; moreover, with one recent exception^50^, these methods perform this segmentation independently for each cell. Such local and cell-specific clustering is problematic for low-coverage single-cell sequencing data because RDRs (and BAFs) have high variance in individual cells.

For CHISEL, we developed a *global* clustering approach that *simultaneously* clusters RDR *x_t,i_* and BAF *y_t,i_* values across *all* bins from *all* cells (Fig. 1B). Specifically, CHISEL uses the *k*-means++ algorithm^64^ to identify clusters of bins which share the same allele-specific copy numbers in every cell *i*. This global clustering approach leverages two observations: (1) all bins from a genome occupy a small number of copy-number states, regardless of their genomic position; (2) all cells from a tumor share a common evolutionary history. This approach extends the global clustering that we introduced in HATCHet^30^ for simultaneous analysis of multiple bulk-tumor samples. We select the number of clusters to minimize the unexplained variance given a certain threshold of tolerance, using standard model selection criteria^65^. An important issue that arises in the global clustering is that one cannot directly compare the BAFs across different bins since we do not know whether the BAF of each bin is the proportion of the allele on either haplotype 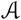 or 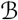. To address this issue, we define the *mirrored BAF ȳ_t,i_* = min{*y_t,i_*, 1 − *y_t,i_*}as in previous studies^20,29^ to guarantee that any pair of bins with similar values of RDRs and mirrored BAFs have the same copy-number state (i.e. the same allele-specific copy numbers).

#### 4.1.3 Inferring allele-specific copy numbers

The third step of CHISEL is to infer a pair {*ĉ_t_, č_t_*}of allele-specific copy numbers for every bin *t* in each cell (Fig. 1C). To do this, one typically needs to know the cell-specific scale factor *γ* that transforms the RDR *x_t_* into the corresponding total copy number *c_t_* = *ĉ_t_ + č_t_*; i.e. *c_t,i_* ≈ *γx_t,i_*. Once one calculates *c_t_*, it is then straightforward to separate *c_t_* into the pair {*ĉ_t_, č_t_*}of allele-specific copy numbers using the BAF *y_t_* since *y_t_* estimates 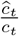 or 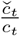. Unfortunately, the scale factor *γ* depends on the genome ploidy *ρ* of the cell, which is generally unknown due to effects of CNAs and WGDs. Existing methods for single-cell sequencing data^6–8,45–50^ infer *γ* using only the RDRs and total copy numbers; for example, Ginkgo^45^ and Cell Ranger DNA^6,48^ minimize the error between the inferred 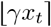 and expected *γx_t_* values of *c_t_* for every bin *t*, i.e. 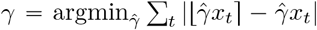. However this approach has two main issues. First, there are generally many equally-plausible solutions for *γ* because *γ* depends on 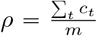 which in turn depends on the total copy numbers to be inferred. Current methods^6,45,48^ use restrictive or biased assumptions on the values of *γ*. Second, because current methods do not consider BAFs, the chosen value of *γ* may result in total copy numbers that contradict the underlying allelic balance, e.g. a total copy number *c_t_* = 1 for a bin *t* with BAF *y_t_* = 0.5.

CHISEL jointly infers the scale factor *γ* and the allele-specific copy numbers {*ĉ_t_, č_t_*} of every bin *t* in a two-stage procedure that integrates both RDRs and BAFs. First, we identify candidate values of *γ* under the assumption that the genome of every cell contains a reasonable number of *balanced* bins, i.e. bins with equal copy numbers *ĉ_t_* = *č* of both alleles. This assumption follows from the observation that bins unaffected by CNAs in a cell have allelespecific copy numbers {1, 1} without WGD, {2, 2} with one WGD, and so on. CHISEL identifies these bins as the largest cluster among the clusters inferred in the second step (Section 4.1.2) whose BAF is approximately equal to 0.5. Second, CHISEL chooses the value *γ* among the candidates and the corresponding pair {*ĉ_t_, č_t_*}of allele-specific copy numbers for every bin *t* using the Bayesian information criterion (BIC) to select among models of varying complexity. Specifically, given a candidate value *γ* and allele-specific copy numbers {*Ĉ_s_, Č_s_*} for all the bins in a cluster *s*, we model the observed RDR *x_t_* and the mirrored BAF *ȳ_t_* of bin *t* ∈ *s* as observations from the following two normal distributions

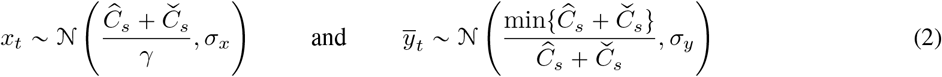

where the sample variances *σ_x_, σ_y_* are estimated from the inferred clusters. For every candidate value of *γ* we find the maximum likelihood estimates for {*Ĉ_s_, Č_s_*} using an exhaustive search, which is feasible as the number of candidate values of *γ* (e.g. 3 when considering the occurrence of at most 2 WGDs) and the number of distinct pairs {*Ĉ_s_, Č_s_*} of allele-specific copy numbers for a cluster *s* are relatively small. Higher values of *γ* always have higher likelihood but also higher model complexity, as they induce more combinations of allele-specific copy numbers; thus we choose the candidate value of *γ* with minimum BIC. Further details of this procedure are in Supplementary Note C.4.

#### 4.1.4 Inferring haplotype-specific copy numbers

The fourth step of CHISEL is to infer haplotype-specific copy numbers (**a**_*i*_, **b**_*i*_) for every cell *i* (Fig. 1D). The challenge is that given the pair {*ĉ_t,i_, č_t,i_*}of allele-specific copy numbers for every bin *t*, we do not know whether *a_t,i_* = *ĉ_t,i_* and *b_t,i_* = *č_t,i_*, or vice versa. The reason for this unknown phasing is that the BAF *y_t,i_* indicates whether the copy number 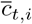 of the minor allele 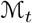 is equal to *ĉ_t,i_* or *č_t,i_*, but we do not know whether 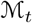 is located on either haplotype 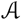 or 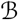. A naive approach that assigns the 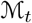 of every bin *t* to the same haplotype generally leads to unlikely scenarios as 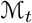 may be determined by different subpopulations of cells in different genomic regions (Fig. S15). While it is impossible to determine the correct phasing given only one sample from a tumor, Jamal-Hanjani et al.^51^ recently showed how to infer haplotype-specific copy numbers in some cases when given multiple bulk samples from a tumor. However the approach in Jamal-Hanjani et al.^51^ has three main limitations that prevent its applicability on single-cell DNA sequencing data. First, the approach relies on the BAFs computed at individual SNPs, making it unfeasible for low-coverage single-cell DNA sequencing data. Second, the approach only determines the presence of different haplotype-specific copy numbers for a specific genomic region but does not phase neighboring regions on the same chromosome. Third, the approach is only successful when different haplotype-specific CNAs are clearly present in *different* samples. We overcome these limitations in CHISEL and infer haplotype-specific copy numbers (**a**_*i*_,**b**_*i*_) jointly across all cells.

The key idea of CHISEL is to phase the minor allele 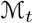 of every bin *t* to the haplotype that minimizes the number of CNAs required to explain the resulting haplotype-specific copy numbers (*a_t,i_, b_t,i_*) across all cells. Specifically, we define the phase *H_t_* of a bin *t* such that 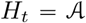 when 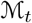 is located on haplotype 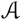 and 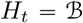 otherwise. Given the phase *H_t_* of bin *t*, we can compute the corresponding haplotype-specific copy numbers (*a_t,i_, b_t,i_*) in every cell 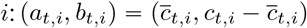 when 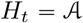 and 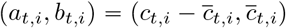 when 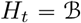. Note that we can easily determine 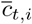 from the BAF *y_t,i_* and the allele-specific copy numbers {*ĉ_t,i_, č_t,i_*}: assuming without loss of generality that *č_t,i_* ⩽ *ĉ_t,i_* we have that 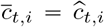 if *y_t,i_* ⩾ 0.5 and 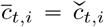 otherwise. To count the number of CNAs that explain a phasing, CHISEL uses the model of *interval events*^17–19^ that model CNAs as events that either increase or decrease the copy numbers of neighboring genomic regions on the same haplotype.

We compute the total number 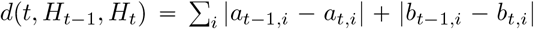 of interval events given the phases *H*_*t*−1_, *H_t_*, as each interval event introduces a difference between *a*_*t*−1_, *a_t_* or *b*_*t*−1_, *b_t_* of two neighboring bins *t* − 1 and *t*. Thus, we seek the phases 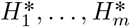 which minimize the total number of interval events across all bins, i.e. 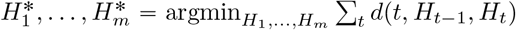. We design a dynamic-programming algorithm to solve this problem based on the following recurrence to compute the minimum number 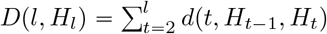 of interval events for the first *l* bins given the phase *H_l_*:

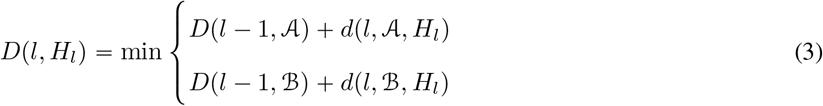

Further details and the proof of correctness are in Supplementary Note C.5.

#### 4.1.5 Inferring tumor clones

The fifth step of CHISEL is to infer distinct subpopulations of cells, or *clones*, with the same complement of CNAs (Fig. 1E). CHISEL uses standard hierarchical clustering to group cells according to their inferred haplotype-specific copy number profiles. More specifically, we define the distance between two cells as the fraction of the genome with different haplotype-specific copy numbers and a threshold is used to determine the maximum difference between cells in the same clone. Next, CHISEL selects the groups of cells that correspond to clones using a minimum threshold on the number of included cells, since we expect that groups composed only of few cells are likely due to noise in the data or errors in the measurements. Further details are in Supplementary Note C.6.

### 4.2 Analysis of breast cancer datasets

#### 4.2.1 Single-cell DNA sequencing data of breast cancer

We analyzed sequencing data from the 10X Chromium Single Cell CNV Solution from 10 tumor datasets of 2 breast cancer patients: (1) 5 adjacent sections from a triple negative ductal carcinoma in situ (patient S0); (2) 3 and 2 technical replicates from two samples of a stage 1 infiltrating ductal carcinoma (patient S1). Each section includes ≈1400-2300 individual cells, whose genome has been sequenced with a sequencing coverage of 0.02-0.05X per each cell. The details of the sequencing procedure are available in the Application Note “Assessing Tumor Heterogeneity with Single Cell CNV” at the 10X Genomics website^5^. The sequencing reads from each dataset were aligned to the human reference genome (hg38 for S0 and hg19 for S1) using the Cell Ranger DNA pipeline^48^.

#### 4.2.2 Inference of allele and haplotype-specific copy number aberrations using CHISEL

We applied CHISEL to analyze every single-cell dataset from patients S0 and S1. We selected only those cell barcodes with a sufficient number of sequencing reads using standard approaches for 10X Genomics data (Supplementary Note C.7). In addition to a BAM file, CHISEL requires two other sources of information: a matched-normal sample and a haplotype phasing for heterozygous germline SNPs. For patient S0 we used section A as a matched-normal sample, as in previous total copy number analysis^5^, because this section contains mostly diploid cells (>91%), which we assumed are normal (non-cancerous cells). For patient S1, we used the available matched-normal sample. In case of a missing matched-normal sample, CHISEL includes an accurate procedure to identify normal diploid cells and to generate a corresponding pseudo-bulk sample (Supplementary Note B.2). We used BCFtools^66^ to identify germline heterozygous SNPs from the matched-normal sample of each patient and we used Eagle2^52^ through the Michigan Imputation Server^67^ to phase germline SNPs with respect to the reference panel HRC (Version r1.1 2016) comprising 64 976 haplotypes^68^. As the HRC panel currently supports the human reference genome hg19 but not hg38, we used LiftoverVcf from Picard tools^69^ to convert the genomic coordinates between the different required builds of the reference genome. For each dataset, we applied CHISEL using the default parameters of haplotype blocks of 50kb and genomic bins of 5Mb. We also applied CHISEL to jointly analyze all the cells of patient S0.

#### 4.2.3 Reconstruction of copy-number trees

We built *copy-number trees* that describe the evolution of the clones identified in section E of patient S0 and the clones identified in joint analysis of all cells of patient S0. A copy number tree has 3 main features: (1) the root corresponds to the diploid clone, (2) the leaves correspond to the other *N* − 1 identified clones, and (3) every branch is labelled by copy number events, with each event either increasing or decreasing the copy numbers of a genomic segment from the parent to the child. We used the model of interval events for CNAs^17–19^ and reconstructed the most parsimonious copynumber tree with the minimum number of events using the haplotype-specific copy numbers inferred by CHISEL. To perform the reconstruction, we separated the two haplotypes of each chromosome and we classified the copy-number events events according to the identified WGD as deletions after WGD (i.e. *del*), deletions before WGD (i.e. *loh*), and duplications before and after WGD (i.e. *dup*).

We used the same approach to identify the events labeling the branches of the copy-number tree that was described in the previous^5^ total copy number analysis for all cells of patient S0. More specifically, we fixed the topology of the tree to be the one reported in previous analysis^5^ and we represented the events as changes in the total copy numbers. In both cases, we ignored small CNAs only affecting few genomic bins.

#### 4.2.4 Analysis of somatic single-nucleotide variants

We pooled the sequencing reads from all cells in each section of patient S0 and ran Varscan^54^ to identify singlenucleotide variants (SNVs). To identify SNVs present in small clones (<100 cells), we relaxed the default parameters of Varscan, selecting the highest confidence 49 356 somatic SNVs with at least 2 supporting sequencing reads from the 8 324 cells in the clones of CHISEL (Supplementary Note C.9). We used SAMtools^70^ to assign each variant read to the corresponding cell through the related barcode. Among all SNVs, 10 551 SNVs were present only in the tumor clones of CHISEL, 541 SNVs present only in the diploid clone, and 38 208 SNVs were in both the diploid clone and the tumor clones. Note that the latter class of mutations likely consist of germline SNPs that were incorrectly classified by VarScan as somatic, false positive variant calls, and early somatic mutations that preceded tumor aneuploidy.

We examined the correspondence between all the identified SNVs and the copy-number tree in two steps. First, we say that a SNV *supports* a branch in the tree if all the cells with the SNV are contained in clones of the subtree descended from the branch. Second, we counted the number of SNVs supporting each non-truncal branch and we assessed whether this number is higher than expected by chance using a permutation test with 10^5^ randomly sampled subsets of cells, each subset containing the same number of cells as in the clones of the corresponding subtree.

We examined the relationship between the variant-allele frequency (VAF) of each SNV and the clonal status of the SNV induced by the CHISEL tree for the 10 551 SNVs identified in the tumor clones. We calculated the VAF of each SNV using the standard definition as the fraction of variant reads over the total number of reads covering the SNV locus. We also defined a *restricted VAF* for an SNV with respect to a subpopulation of cells by restricting to sequencing reads with barcodes matching the cells in the subpopulation. In particular, we computed a *left-restricted VAF* and a *right-restricted VAF* by restricting to the sequencing reads from cells belonging to the two main branches of the CHISEL tree. Next, we classified the SNVs according to the CHISEL tree by separating SNVs into *clonal* SNVs, which are present in all tumor clones, and *subclonal* SNVs which are unique to either the left or right branch.

Distinguishing true positive from false positive SNVs is complicated in this dataset due to the low number of variant reads for many SNVs. Thus, we restricted attention to high prevalance SNVs that were present in multiple clones of the same branch, resulting in 594 SNVs unique to the left branch, 1 632 SNVs unique to the right branch, and 2 798 clonal SNVs present in both branches. The remaining low-prevalance SNVs have both low VAFs (Fig. S13) and low restricted VAFs (Fig. 4C) in both branches, underscoring the low confidence in these mutation calls. Further details of the VAF analysis are in Supplementary Note C.10.

## Acknowledgments

We thank Lance Hepler and Karthik Ganapathy from 10X Genomics for providing additional data for our study, for providing access to the published data of the total copy number analysis, and for the useful feedback. This work is supported by a US National Institutes of Health (NIH) grants R01HG007069 and U24CA211000, US National Science Foundation (NSF) CAREER Award (CCF-1053753) and Chan Zuckerberg Initiative DAF grants 2018-182608 to BJR.

## Code availability

CHISEL is available on GitHub at https://github.com/raphael-group/chisel.

## Data availability

All the 10x Genomics Chromium BAM files used for the 5 sections of patient S0 have been downloaded and are publicly available at https://www.10xgenomics.com/resources/datasets/. In particular:

1. Section A: https://support.10xgenomics.com/single-cell-dna/datasets/1.0.0/breast_tissue_A_2k
2. Section B: https://support.10xgenomics.com/single-cell-dna/datasets/1.0.0/breast_tissue_B_2k
3. Section C: https://support.10xgenomics.com/single-cell-dna/datasets/1.0.0/breast_tissue_C_2k
4. Section D: https://support.10xgenomics.com/single-cell-dna/datasets/1.0.0/breast_tissue_D_2k
5. Section E: https://support.10xgenomics.com/single-cell-dna/datasets/1.0.0/breast_tissue_E_2k

The 10x Genomics Chromium BAM files used for the 5 datasets of patient S1 are pending release. All the processed data for all sections of patient S0 and S1, as well as all the results of CHISEL are available on GitHub at https://github.com/raphael-group/chisel-data.

